# Paying attention to natural scenes in area V1

**DOI:** 10.1101/2023.03.21.533636

**Authors:** Andreea Lazar, Liane Klein, Johanna Klon-Lipok, Mihály Bányai, Gergő Orbán, Wolf Singer

## Abstract

Natural scene responses in the primary visual cortex are modulated simultaneously by attention and by contextual signals about scene statistics stored across the connectivity of the visual processing hierarchy. Here, we hypothesized that attentional and contextual top-down signals interact in V1, in a manner that primarily benefits the representation of natural visual stimuli, rich in high-order statistical structure. Recording from two macaques engaged in a spatial attention task, we found that attention enhanced the decodability of stimulus identity from population responses evoked by natural scenes but, critically, not by synthetic stimuli in which higher-order statistical regularities were eliminated. Population analysis revealed that neuronal responses converged to a low dimensional subspace for natural but not for synthetic images. Critically, we determined that the attentional enhancement in stimulus decodability was captured by the dominant low dimensional subspace, suggesting an alignment between the attentional and natural stimulus variance. The alignment was pronounced for late evoked responses but not for early transient responses of V1 neurons, supporting the notion that top-down feedback was required. We argue that attention and perception share top-down pathways, which mediate hierarchical interactions optimized for natural vision.

## Introduction

Neuronal circuits across the visual hierarchy make efficient use of limited resources to encode complex natural scenes by exploiting their structural regularities (1, 2). Growing evidence suggests that during perceptual inference the visual system employs a hierarchical internal model of the visual environment that integrates current sensory evidence with previously acquired knowledge of natural scene statistics (3, 4).

When visual attention is directed towards a specific spatial location, it is thought to facilitate the perception of the targeted sensory input by prioritizing its processing. The signatures of this process can already be observed in primary visual cortex (V1). In this area, spatial and object centered attention modulate the firing rates of neurons responding to the selected stimuli (5–9). Responses in V1 are also shaped by context-dependent top-down signals that convey information stored in the hierarchically organized internal model of the world, required for the parsing of visual scenes (4). Thus, attentional mechanisms must interact with other sources of top-down influences raising the key question: how do these two processes — one that reflects the current allocation of attention and the other, the stored knowledge (priors) about statistical regularities of natural environments — cooperate in modulating responses in V1.

Here we performed parallel recordings of neuronal responses in the primary visual cortex of two macaque monkeys engaged in an attention task. In this paradigm, attention was directed to one of two identical images by a spatial cue that was presented delayed from the stimulus onset. By measuring multiunit activity of a population of neurons with receptive fields overlapping with one of the images we tracked the contribution of the top-down effect of attention on the distributed stimulus representation. We found convincing enhancements in natural scene encoding with attention and hypothesized that these enhancements rely on refinements of the responses to high-level features present in natural images, such as regularities in spatial, contrast and frequency structures or texture properties. To test this hypothesis, we independently manipulated a second form of top-down modulation, arising from the statistical structure of the input images. We constructed synthetic stimuli that lacked the higher-level features characteristic of natural scenes and found that, for these stimuli, the attentional benefits in stimulus encoding vanished; however they could be recovered when the synthetic stimuli were modified to contain structured contours. To understand how attentional modulations might interact with the representation of stimuli, we investigated the geometry of population responses using a combination of dimension reduction methods and decoding analysis. Our analysis revealed that the evoked neuronal population responses to natural scenes converged to a compact low dimensional representation, which coincided with the subspace where the dominant attentional signal could be identified.

## Results

We obtained parallel multisite recordings of multi-unit activity (MUA) (example trials in Figure 1A) and local field potentials (LFP) (32 channels, chronically implanted microdrive, Figure 1B) from area V1 of two awake behaving macaques (*Macaca mulatta*), while they performed a spatial attention task. Trials were initiated by a lever press while the monkey maintained fixation. After 500 ms two identical stimuli were presented at symmetrical locations on either side of the vertical meridian (distance from fixation spot 2.3-3.2° of visual angle), one of which covered the receptive fields (RFs) of the recorded units (example of RF centers in the two monkeys in Figure 1C). The fixation spot changed color, 700 ms after stimulus onset, cueing the monkey towards the “target” stimulus. Monkeys were rewarded if they responded to the rotation of the target stimulus. If they responded to the rotation of the non-cued stimulus, the distractor, reward was withheld and the trial was aborted. We performed analysis in 200 ms long sliding windows for the response interval from stimulus onset to 1900 ms (stimulus-aligned data), and for the 500 ms interval preceding the stimulus change (change-aligned data, time of change (TC)), thus excluding any activity evoked by the stimulus rotation or the motor response (further details in Materials and Methods).

**Fig. 1.**
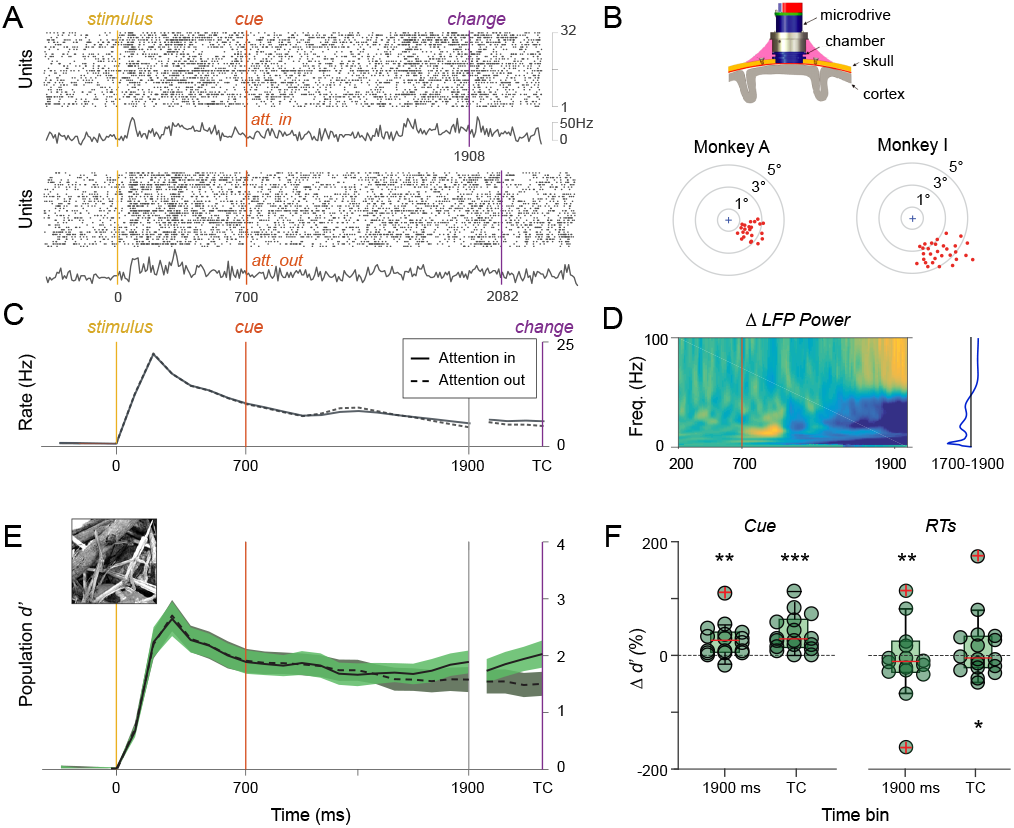
Neuronal population recordings from primary visual cortex during an attention task with natural scene stimuli. (A) Multi-unit responses to the same stimulus in two example trials, corresponding to the attention in and attention out conditions. Colored lines represent timing of stimulus onset (yellow), attention cue (red) and stimulus change (purple). (B) Top: Chronic implantation device with movable electrodes (GrayMatter probe). Bottom: Receptive field locations of multiunits (dots) in the two monkeys. Cross indicates the fixation point. (C) Grand average firing rates in the attention-in and attention-out conditions (colored lines as in A). Spectrogram of the change of LFP power between the attention-in and attention-out conditions. A gradual shift in power from low to high frequencies starts *∼*300ms after the attentional cue (marked by vertical line) (E) Effects of attention on stimulus discriminability along the trial, quantified by population discriminability (d’). (F) Effects of attention on stimulus discriminability shown across sessions (reported in % change) for the late stimulus responses (data aligned on stimulus onset in the 1700–1900 ms window) or stimulus change (200 ms window before change). Left panel: trials are separated based on cue information. Right panel: trials from the attention-in condition are sorted based on reaction time and separated in two halves. y-axis shows percent increase in *d*′ for *in* vs *out* trials (Cue) or *fast* vs *slow* trials (RTs).

We found that attention increased the average firing rates only modestly in V1, and the changes were significant only for the change-aligned and not the stimulus aligned data (Figure 1C and S1, Wilcoxon signed-rank test; [1700-1900ms], P = 0.1330 n.s.; [TC-200, TC], P = 0.0279; n = 18 sessions in 2 monkeys). In spite of the small firing rate changes, we observed a strong shift in LFP power from low to high frequency oscillations with attention, which started *∼* 300ms after the cue and increased gradually with the expectancy of the stimulus change (Figure 1D, individual animals in S4). Across trials, reaction times in the attention-in condition were positively correlated with the LFP power in the theta and beta bands and negatively correlated with the LFP power in the gamma band (Spearman correlation; theta *r* = 0.18, *p* = 1*e* − 11 ; beta *r* = 0.08, *p* = 0.002 ; gamma *r* = −0.12, *p* = 1*e* − 05), suggesting a strong task-related effect.

Our initial goal was to examine the impact of attention on the stimulus specificity of the sustained population responses to natural scenes. Few electrophysiology studies have focused on the representation of natural stimuli in primary visual cortex (10–12), and how attention affects the decodability of natural scenes in V1 is still unclear. We quantified the stimulus-specificity of V1 neuronal population responses by measuring the differences in spiking patterns evoked by different stimuli compared to their variability across trials (population *d*′, Materials and Methods). We found a significant increase in population *d*′ with attention, i.e. responses evoked by different stimuli were more differentiable when the stimuli had been cued (attention-in) than when they had not (attention-out), particularly in the time windows preceding the stimulus change (Figure 1C and D, Wilcoxon signed-rank test; [1700-1900ms], P = 0.0012; [TC-200, TC], P = 0.00023; n = 18 sessions in 2 monkeys). These attentional benefits in natural scene discriminability were significant in individual animals and occurred both in the presence (monkey A), and absence (monkey I) of average firing rate changes (Figure S2). Importantly, a significant difference in stimulus discriminability, was not only elicited by the cue, but also by intrinsic attention-related factors: when the trials from the *attention-in* condition were sorted based on the monkeys’ reaction times (RTs), we found that trials with faster RTs had higher *d*′ values compared to those with slower RTs (Figure 1F, Wilcoxon signed-rank test; [1700-1900ms], P = 0.0043 ; [TC-200, TC], P = 0.0123; n = 18 sessions in 2 monkeys). The relationship between *d*′ and RTs points to a graded improvement in stimulus decodability proportionate to the intensity of the attentional allocation, suggesting a link between the modulatory signals identified in V1 and behavior.

Response properties of neurons at higher levels of the processing hierarchy are known to capture the higher-order regularities of natural scenes (13). Indeed, when gauging V2 activity by using natural texture images, specific removal of high-level statistics was shown to result in dampened responses of V2 neurons (14). Given the hierarchical and reciprocally connected structure of the visual cortex, information extracted and encoded at higher levels of the processing hierarchy can constrain the activity at lower levels of processing through feedback. Consequently, such contextual feedback modulation is likely more effective when natural highlevel features are present in stimuli (4). We hypothesized that if contextual modulation and attention share top-down pathways then the presence of high-level features constitutes a prerequisite for attentional enhancements in stimulus encoding across neuronal populations in V1.

To test this hypothesis we matched the natural scene stimuli, in every recording session, by an equal number of synthetic control images. The synthetic images were constructed in two ways. First, by filter-scrambling, to remove spatial correlations between low-level features (Figure 2A, details in Materials and Methods), which was previously shown to reduce top-down influences on V1 responses (4). Second, by phase-scrambling, to remove high-level regularities from images while leaving the spectral content intact (Figure 2B), which was previously shown to reduce both the intensity and specificity of responses of V2 neurons (14). By construction, the synthetic controls contained no high-level features, but retained basic image properties: either the low-level structure preferred by V1 cells or the spectral content of the original natural scenes.

**Fig. 2.**
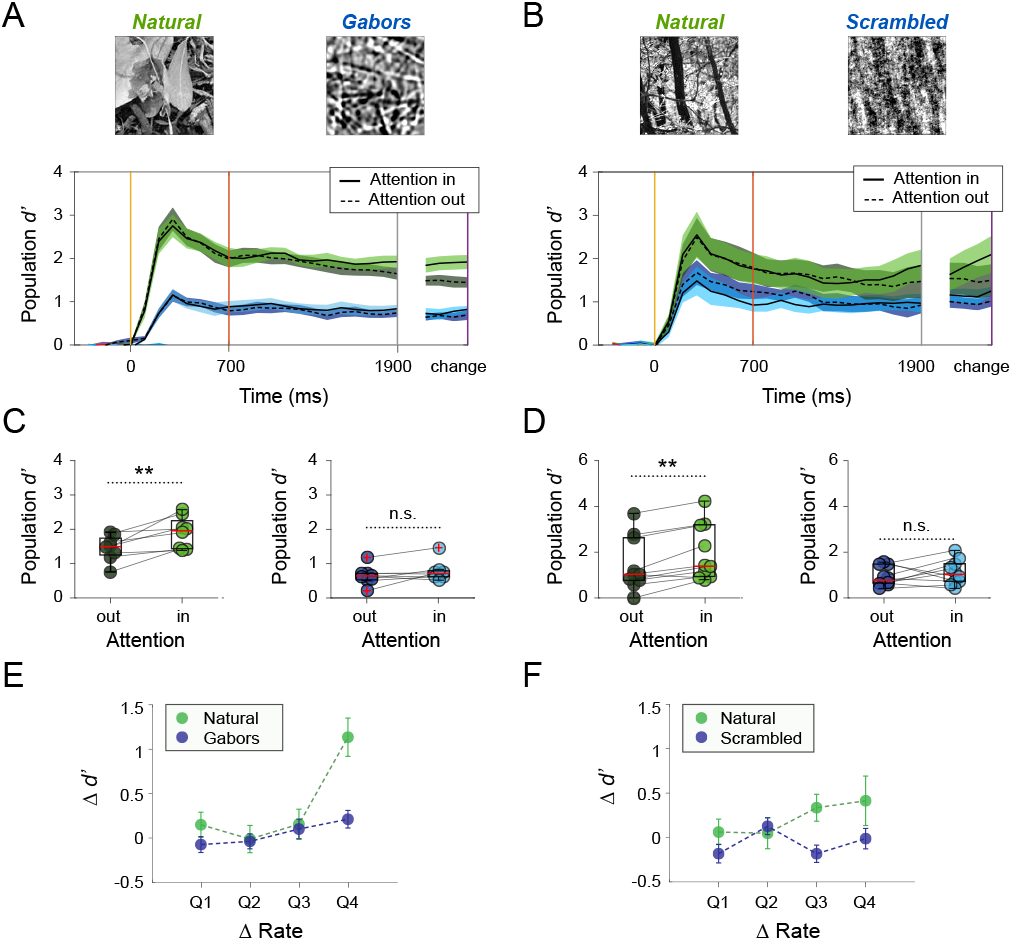
Attentional modulation depends on stimulus content. (A) Stimulus discriminability *d*′ for natural scenes (green) and synthetic images composed of independent Gabor filters (blue), shown along the trial, 200 ms spike-count vectors, 100 ms sliding resolution. (B) Stimulus discriminability *d*′ for natural scenes (green) and phase-scrambled images (blue). (C,D) Attentional modulation of *d*′ for natural (green) and synthetic images (blue) in the time window preceding the stimulus change (circles depict individual sessions). (E,F) Change in *d*′ as a function of change in firing rate with attention. Analysis performed across all recorded units. Responses were sorted by amplitude of rate change and grouped into quartiles (marked on *x*-axis). Large increases in firing rate with attention result in larger positive changes in *d*′ for natural (green), compared to synthetic stimuli (blue). Error bars indicate the standard error of the mean.

The synthetic controls matched the natural scenes in luminance contrast and evoked, on average, similar firing rate responses across the recorded neuronal populations in V1 (Figure S3). In addition, the synthetic stimuli observed a task-related shift in LFP power from low to high frequencies with attention, of similar magnitude to that observed for natural stimuli (Figure S4, compare dotted and continuous lines in subplots B and F). Reaction times across trials in the attention-in condition for synthetic stimuli were positively correlated with the LFP power in the theta and beta bands and negatively correlated with the LFP power in the gamma band (Spearman correlation; theta *r* = 0.11, *p* = 3*e* − 5 ; beta *r* = 0.07, *p* = 0.008 ; gamma *r* = −0.11, *p* = 3*e* − 04) and were indistinguishable from reaction times to natural stimuli (Wilcoxon signed-rank test; P > 0.3, n.s.; n = 18 sessions in 2 monkeys), suggesting a similar level of engagement. One notable difference was that the LFP power in the gamma band was higher for natural scenes compared to synthetic images (Figure S4), consistent with previous suggestions that visual stimuli with higher structural predictability result in stronger gamma oscillations (12). Nonetheless, attentional modulation of LFP power in the gamma band showed a reduction in one monkey and a slight increase in the other, and was therefore difficult to interpret (Figure S4).

In spite of the overall similarities between responses to natural and synthetic stimuli, in both overall firing amplitudes and task-related LFP dynamics, the spike-count vectors evoked by synthetic stimuli were considerably less discriminable than those evoked by natural scenes in the time interval preceding the stimulus change (Figure 2A and B). Most importantly, in agreement with our hypothesis, the discriminability of neither type of synthetic stimuli was enhanced by attention (Figure 2C and D, Wilcoxon signed-rank test; [TC-200, TC]; natural images P = 0.0078, Gabors P = 0.25 n.s.; n = 8 sessions in 2 monkeys; natural images P = 0.0039, scrambled P = 0.375 n.s.; n = 10 sessions in 2 monkeys). Moreover, when responses of all recorded units were pulled together and sorted by amplitude of rate change with attention and grouped into quartiles, we found that equally strong increases in firing rates resulted in *d*^*/*^ enhancements for natural but not for synthetic stimuli (Figure 2E, 256 units; and F, 315 units). Thus, while the grand average discharge rates were similarly modulated for natural and synthetic stimuli, the attentional enhancement in stimulus discriminability was specific to natural scenes, suggesting a dissociation between the attentional modulation of firing responses and the resulting gains in stimulus specificity (Figure S3).

These findings raise the question why responses to natural scenes profit more from attentional refinement than responses to manipulated stimuli or, in other words, what is special about natural scenes? Previous research has shown that natural images are statistically redundant, since light-intensities at neighboring locations are likely to be correlated and consequently, they can be efficiently compressed (15, 16). Structured compressible visual stimuli are well captured by neuronal population dynamics in low-dimensional manifolds (12, 17–19), but see (20). How could such low-dimensional collective representations of natural images in V1 be further optimized by the allocation of top-down attention?

Given that, in the current task, the stimuli precede the attentional cue, we could directly enquire whether the variance added by attention was orthogonal to, or belonged to the same dimensions as the variance produced by the stimulus. Specifically, in each recording session, we projected the activity from the time window preceding the stimulus change (1700-1900 ms) into the principal component space defined by activity recorded after the stimulus onset but before the presentation of the attentional cue (500-700 ms). We reasoned that if the attentional variance was largely aligned to the stimulus variance, the attentional differences in stimulus decodability would become apparent in the low-dimensional space described by the first components. This is indeed what we found (examples in Figures 3A and S5).

**Fig. 3.**
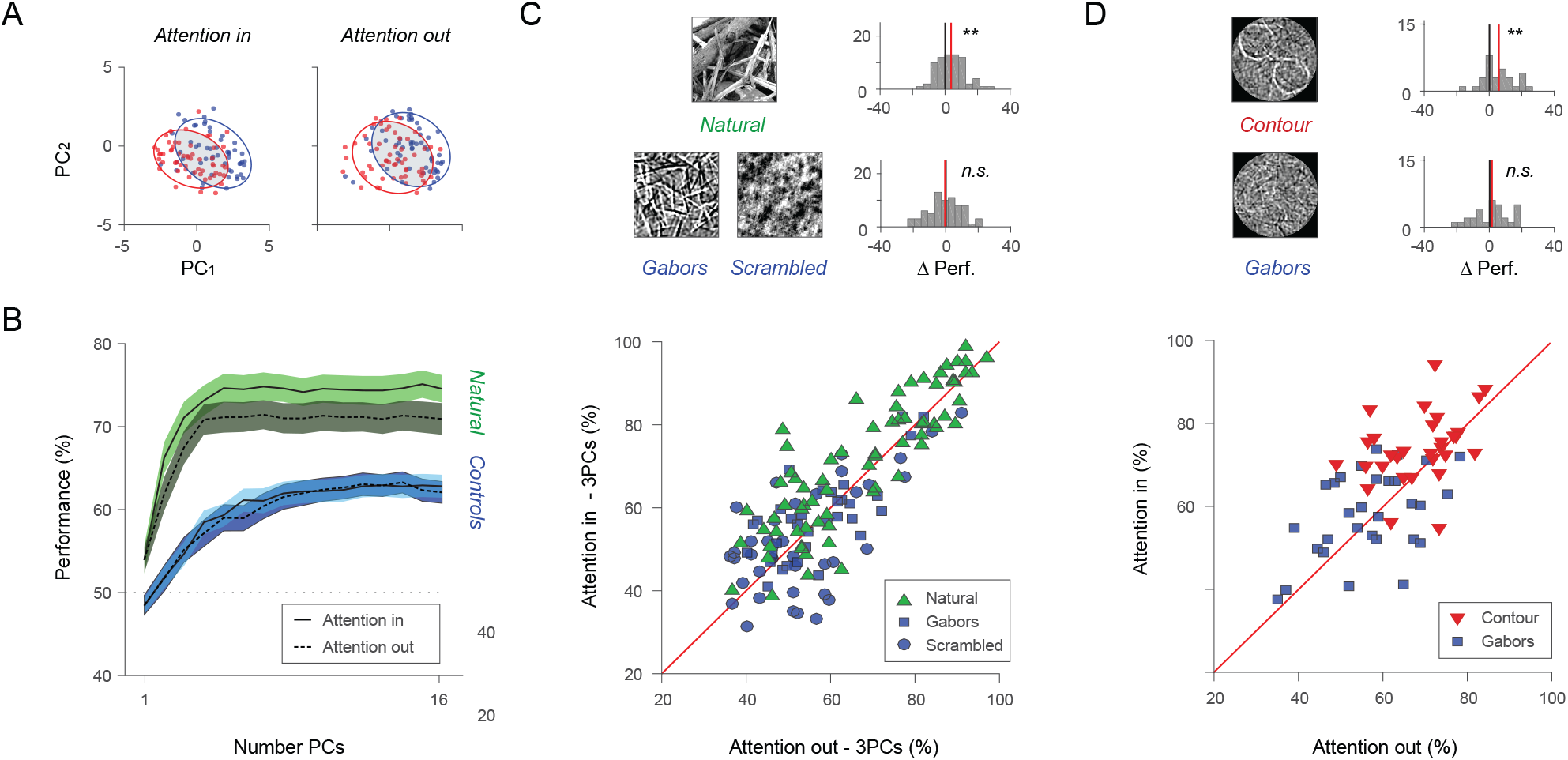
Effects of attention on stimulus encoding in principal component space. (A) Example session depicting population responses to two natural scene stimuli (red and blue) for the two attentional conditions in the space described by the first 2 principal components (200 ms spike-count vectors, 1700-1900 ms from stimulus onset; each point represents a trial). (B) Stimulus decoding performance in principal component space for natural scenes (green) was higher than for synthetic stimuli (blue) and was modulated by attention (based on 1700-1900 ms spike count vectors; PCA was performed on pre-cue activity 500-700 ms window). The shaded areas indicate the standard error of the mean. Attentional differences in stimulus decoding are apparent from a low number of PCs, suggesting an alignment between attentional and stimulus variance. (C) Impact of attention on performance scores in low-dimensional projections (first 3 PCs) depends on stimulus type. Scatterplot shows performance scores for natural scenes (green) and synthetic controls (Gabors, blue squares; scrambled, blue circles; markers represent stimulus pairs n = 71; 18 recording sessions, 2 monkeys). Differences in performance with attention are significant for natural stimuli (top histogram, ** p-val < 0.01) but not controls (bottom histogram, n.s. p-val > 0.05). (D) Contour stimuli are compared to synthetic controls (n = 30 stimulus pairs from 5 recording sessions in 1 monkey). Differences in performance with attention are significant for contour stimuli not controls.

We quantified the attentional differences on natural-scene representations in principal component space by applying a decoding technique. Specifically, a cross-validated Bayesian decoder was trained to predict stimulus identity based on data projections (spike-count vectors over the 1700-1900 ms interval) into the pre-cue PCA space described by the first *k* principal components, and test performance was estimated in this same space based on unseen trials (Figure 3B, 5-fold validation, details in Materials and Methods). We found that a small number of components captured the majority of variance produced by natural stimuli before the onset of the cue, allowing natural scenes to be well distinguished in principal component space (Figure 3B, decoding performance range 61.8 − 75.13% for *k* >=2). Importantly, the same components captured a large fraction of the attentional effects, as reflected by the significant modulation of population responses in low-dimensional subspaces (Figure 3B, Wilcoxon signed-rank test; P < 0.015 for *k* >=2; Holm-Bonferroni correction showed significance for all *k* >=2; n = 71 stimulus pairs, 18 sessions, 2 monkeys; scatterplot and histogram of attentional effects for *k* = 3 are shown in Figure 3C). In comparison, the synthetic images performed more poorly (Figure 3B, decoding performance range 51.7 − 63.3% for *k* >=2) and showed no attentional modulation (Figure 3B and C, Wilcoxon signed-rank test P > 0.05 for all *k*; n = 71 stimulus pairs, 18 sessions, 2 monkeys), in spite of residing in principal component spaces with similar levels of overall variance (Figure S6B).

In a final set of experiments, we generated synthetic images that combined Gabor functions into simple contour-like patterns, thus introducing the kind of higher-level structure expected to elicit differential responses at higher processing stages. In these additional datasets, both main effects described previously were reproduced: the structured contour stimuli were well distinguished in principal component space, while the unstructured controls were not (decoding performance range contour stimuli 66.4 − 74.7% and synthetic Gabors 54.9 − 60.4% for *k* >=2, Figure S6) and the attentional effects were specific to the contour stimuli and captured already by a low number of components (contour stimuli Wilcoxon signed-rank test; P < 0.05 for all *k* >=2; n = 30 stimulus pairs, in 1 monkey; control images P>0.05 for all *k*, Figure S6; scatterplot and histogram of attentional effects for *k* = 3 Figure 3C). Interestingly, trial-shuffling within stimulus condition reduced the attentional differences in decoding performance for the contour stimuli and the original natural scenes, suggesting that in these low-dimensional projections decoders benefitted from the intact correlation structure present in the data (Figure S6). Overall, by constructing synthetic images with controlled statistical structure, we confirmed that the attentional benefits in stimulus encoding across collective neuronal responses in V1 were specific to images containing higher level structural regularities. Such images are more likely to engage structured feedback from higher-levels of processing.

Critically, we found that the dimensions that captured the majority of variance produced by natural stimuli also captured a large fraction of the attentional effects. To control for the specificity of this alignment, we constructed three alternative projections of the same spike-count vectors preceding stimulus change (1700-1900ms) and assessed how these spaces captured both the natural stimulus and the attention signal. We considered a random orthogonal basis (Figure 4A, yellow), a PC space constructed based on pre-stimulus spontaneous activity (Figure 4A, orange) and a PC space based on early evoked activity (Figure 4A, purple). The minimum number of principal components required to reach 90% of peak performance accuracy for the attention in condition was significantly higher for the three projections compared to the original pre-cue projection (Figure 4B, compare yellow, orange, purple to green; Wilcoxon signed-rank test, P<0.001). In addition, all three alternative projections yielded weaker attentional differences for both the first three and five components (Figure 4C; Wilcoxon signed-rank test, P<0.01). Most interesting was the difference between the projections constructed from the early and late evoked responses, which were compared directly in Figure 4D. Session-by-session comparison of attention-in and attention-out decodability, in these two projection spaces, revealed a dual effect: 1, Decoding of stimulus was more efficient in the basis constructed based on late responses (green) than that based on early responses (purple), irrespective of the attentional state (Wilcoxon signed-rank test, P <0.001 attention-in; P <0.001 attention-out); 2, Attention made a significant contribution to decoding in the basis constructed using late responses (Figure 4D, green histogram; Wilcoxon signed-rank test, P = 0.000014). Note that neither of these two projections had information about which stimulus to attend, since both are precue, therefore it is striking that a precise alignment developed later in the trial. This result indicates that during stimulus presentation the population response is transformed into the space in which attention can also be effectively deployed.

**Fig. 4.**
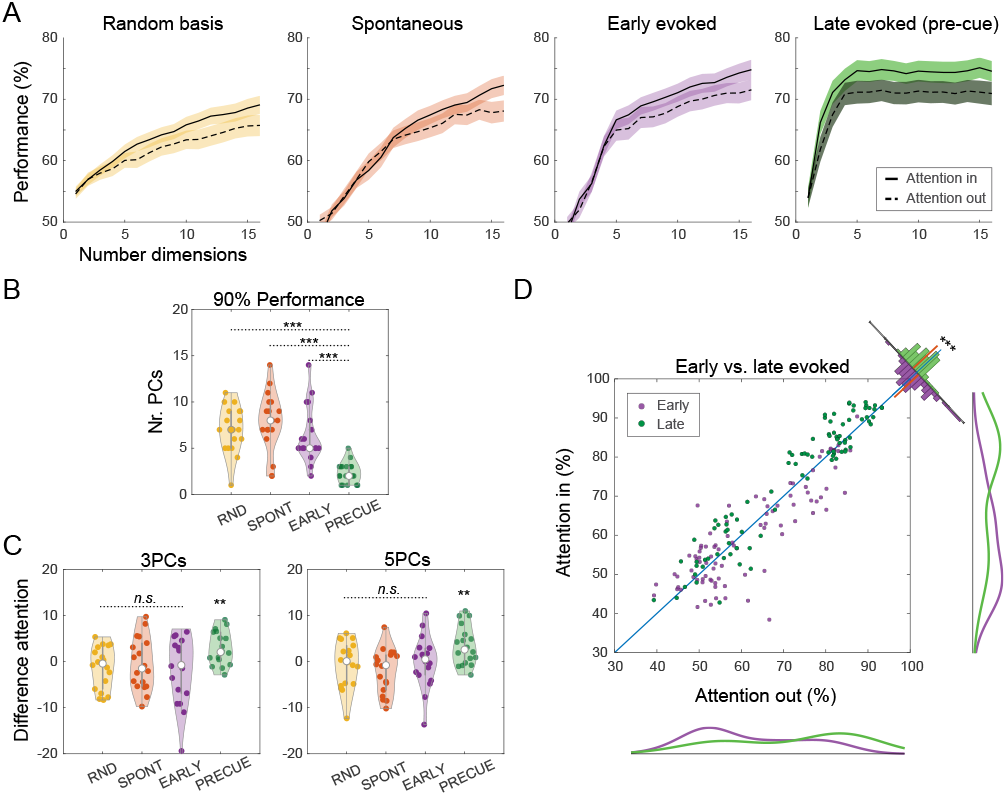
Geometry of population activity in response to natural scenes. (A) Spike-count vectors from the post-cue interval (1700-1900 ms) are projected in four different bases: a random projection space (yellow), a PC-space based on spontaneous activity (orange, interval -200-0 ms), a PC-space based on early evoked activity (purple, 100-300 ms window), and a PC-space based on late evoked activity (green, pre-cue, 500-700 ms window, same as Figure 3B). When using PC decompositions, the dimensions correspond to the first PCs of the PCA basis. Average performance of the stimulus decoder, across all stimulus pairs, from all recording sessions, shown as a function of the number of dimensions used to reconstruct population activity. Attention-in (solid line) and attention-out trials (dashed line) are decoded separately. (B) Number of dimensions necessary to reach 90% of peak decoding accuracy, shown across recording sessions. The late evoked projection space (green), requires fewer than five components. Interestingly, the early evoked projection space (purple) requires up to 15 components to reach the same decoding accuracy, suggesting a higher dimensional encoding space. (C) Attentional difference in decoding accuracy (attention in - attention out), using a three or five-dimensional decoding space, shown across sessions. Differences are significant in the projection space defined based on late evoked activity, but not the alternative projection spaces. (D) Direct comparison of post-cue activity in the early (purple) and late evoked (green) projection spaces. A projection of the post-cue activity is used to decode stimulus identity from attention-in and attention-out trials. Dots indicate individual sessions, data pulled for the first 5 components. Marginals of decoding performances along the axes (solid lines) show overall decoding performance differences for early vs late responses, indicating differences in the dimensionality of the population responses. Difference of attention-in and attention-out decoding performance (histogram perpendicular to the identity line) indicates differences in the alignment of the attentional modulation with the dominant activities in early and late responses. Both stimulus decoding and the attentional benefits are stronger in the PC space constructed on late evoked activity.

## Discussion

In this study, we found that attention improves the representation of natural scenes across neuronal population vectors in area V1. By constructing synthetic stimuli with controlled statistical structure, we could link the attentional benefits in stimulus encoding to the presence of higher-order regularities that are known to be abundant in natural images and are primarily represented at higher-level areas of the ventral stream. Population analysis revealed that the attentional signal was aligned with the compact subspace carrying information about stimulus identity. Temporal evolution of the stimulus-representing subspace revealed that alignment was not present in early responses to natural stimuli, but it emerged later, still preceding the delivery of the attentional modulation. Taken together, attentional enhancement of V1 representation of natural stimuli harnesses high-level statistical structure represented in higher visual cortical areas and is carefully and specifically aligned with the activity carrying information about natural structure.

In our experiments, animals were trained to respond swiftly to the rotation of the cued stimulus, not to recognize particular features in the image. Thus our task was a classical spatial attention task. In agreement with previous studies allocating spatial attention caused moderate increases in discharge rate, reduced power of low frequency oscillations and enhanced power in the high frequency bands of local field potentials. As expected, these effects did not depend on stimulus structure. In agreement with psychophysical and previous electrophysiological investigations this indicates that spatial attention enhances the salience of responses (21). However, allocating spatial attention had the additional effect of enhancing selectively the decodability of population responses of V1 to the attended stimulus, provided that the stimulus contained higher order statistical regularities characteristic of natural scenes. These effects on decodability could not be attributed to a global increase of excitability because they occurred also in the absence of rate changes. Extensive evidence supports the notion that the structural features characteristic of natural scenes are processed at higher levels of the hierarchy. More recently, it has been shown that receptive fields of V1 neurons can be more complex than earlier described (22). It is possible that the receptive fields of the recorded neurons are actually matching the natural scenes better than the synthetic stimuli, which could also explain the higher discriminability of natural scenes. Our experimental design relied on a limited number of images, which prevented us from a full characterization of the response properties of the recorded neurons. However, we designed synthetic stimuli based on evidence that the statistics we manipulated are characteristic to the secondary visual cortex (13, 14, 23), and therefore such manipulations are expected to affect the information that is fed back to V1, even when the targeted responses are complex. Further, the observed enhanced discriminability was not present when using a PCA based on early evoked responses, in contrast with the expectation that a feed-forward pass of information is sufficient to account for the better discrimination of natural-like stimuli. Taken together, these imply that allocation of spatial attention must have interfered with feature sensitive top-down mechanisms, raising the question how these two processes interact.

Visual scenes are evaluated by comparing sensory evidence with previously acquired priors about the statistical structure of natural environments (1). These internal priors are stored in the functional architecture of cortical circuits at all levels of the visual processing hierarchy and some of these circuits get refined by experience to capture characteristic properties of the visual environment (2, 3, 24). Recombination of feedforward connections renders neurons selective for increasingly complex constellations of features (25–27), and the abundant horizontal intra-areal and feed-back connections between processing levels allow for contextual modulation of these feature selective responses (28, 29). These modulations impact stimulus saliency (30) and perceived brightness (31, 32), support perceptual grouping (33), and figure-ground segregation (34–37). The electrophysiological correlates of these interactions consist of changes in discharge rate and/or synchrony and these effects tend to have longer latencies than the initial phasic responses. Therefore it has been concluded that the context sensitive processes are mediated by recurrent interactions within cortical areas and top down signaling across processing stages. In the following we discuss how allocation of spatial attention that is also supposed to be mediated by top-down connections interacts with these feature sensitive mechanisms.

Attentional influences on visual processing have traditionally been divided into spatial (38–40) and object/feature-based attention (41, 42) and it has been proposed that both contribute in complementary ways to the parsing of image content (43, 44). Our results support this notion and provide some indications as to the mechanisms underlying these complex interactions, in the context of natural stimulation. In the present task, the allocation of spatial attention contributed additional variance in the first principal components of responses to natural but not to manipulated images and thereby enhanced decodability of the former. The finding that spatial attention had no effect on decodability of manipulated stimuli indicates that spatial attention has per se no refining effect on distributed stimulus representations in V1, but selectively improves representations of stimuli characterized by the higher order regularities of natural scenes. Abundant evidence indicates (45) that these higher order regularities are evaluated by downstream areas of the visual processing hierarchy. Therefore, the enhanced decodabilty of responses to natural images is likely to have been mediated by top-down signals from these areas. This raises the question, why these high-level processes were more involved when spatial attention was allocated to the stimulus. One possibility is that higher level processes do not engage by default even when stimuli match high-order priors but get involved only for stimuli to which spatial attention is allocated. In this case spatial attention would be a prerequisite for the engagement of mechanisms that provide top-down signals commonly attributed to feature or object specific attention, suggesting some hierarchy in the interactions between spatial and object centered or feature specific attention. An alternative possibility is that higher level processes engage by default when stimuli match high-order priors and work cooperatively alongside spatial attention. In this case, the visual system performs a search that attempts to infer task-relevant features of an image based on both the spatial aspects of the visual scene and the low-level and high-level structural regularities. Thus spatial and object-based attention act in unison and share an internal representation of features, with the inference slowly unfolding over reciprocal interactions across multiple hierarchical levels of processing.

The latter interpretation is supported by the observation that the natural scenes could be discriminated surprisingly well, given the relatively low number of units and their location in area V1 (Figures 2 and 3), regardless of the attentional cue. The fact that the discriminability of natural stimuli was high also when attention was directed away from the stimulus implies that the natural scenes were efficiently encoded, irrespective of the attentional state. Previous studies found that an efficient encoding of global scene statistics remained possible in situations associated with reduced visual attention (46). In such cases, the visual cortex is thought to extract a compressed “summary” code that does not capture the full distribution of local details, yet provides a good representation of group features. Since natural scenes are structured, redundant, low-dimensional images, they are compressible. In comparison, the low-level synthetic images are difficult to compress and must be represented exhaustively, without the help of internally generated or previously acquired priors on summary statistics. Thus the resulting neuronal activity vectors to synthetic stimuli are likely to inhabit higher-dimensional or more variable substates, potentially accounting for the overall poorer performance of the classifier.

Principal component analysis was used to capture the efficient encoding of natural scenes across neuronal populations in V1. In principal component space, the response vectors to natural stimuli reflected their low-dimensionality and could be well described by few principal components. The higher-order stimulus structure, characteristic of natural scenes, was thus well-separated by low-dimensional subspaces. In a sense, these subspaces reflect some of the higherorder selectivity normally associated with responses of individual neurons at higher levels of processing (47, 48). The finding that spatial attention enhanced the encoding of natural scenes along the dominant representational dimensions, suggests refined, cooperative interactions across multiple levels of the visual hierarchy. Earlier, behavioral studies hinted at interactions between attentional effects and regularities beyond the simple features represented in lower visual areas (49). Here, the match between representational and attentional signals in V1 shows a remarkable alignment, likely advantageous for efficient processing. This match appears compatible with previous results highlighting similarities between the effects of representational learning and attention in downstream area V4 (50).

Spatial attention modulated the dynamics of responses as reflected by changes in the frequency distribution of LFP power. A shift in LFP power from low to high frequencies built up gradually from the onset of the cue to the temporal window preceding the stimulus change, for both natural and synthetic stimuli (Figures 1 and S4). This kinetics resembled a hazard function reflecting the increasing probability of having to execute a response, suggesting that the reduced power of the low frequency oscillations was probably related to the increased readiness to act. In agreement with this interpretation the oscillatory power in the theta and beta bands of responses to the target stimulus was positively correlated with reaction times in the attention-in condition. These results are consistent with previous reports. Beta oscillations have been shown to decrease during the preparation of a motor response (51) and theta band power has been shown to decrease with attention (52). The strength of gamma band oscillations was negatively correlated with reaction times in trials corresponding to the attention-in condition, in agreement with previous findings from area V4 (57) and also with our observation that the gradual shift in power from low to high frequency oscillations is related to the readiness to act (see above). In agreement with previous work is also the finding that attention enhances the power of the broad band high frequency activity that likely reflects increases spiking and synaptic activity (58). In addition to these attention dependent effects on dynamics, we observed a build up of gamma oscillations for natural, but not for synthetic stimuli, over the course of the trial (Figure S4). Gamma oscillations result from a feedback loop between pyramidal cells and fastspiking interneurons (53, 54) and are thought to act as an internal resonance filter of stimulus content (12, 55). Synchronisation of discharges in the gamma frequency range increases for responses to features that are well predicted by the embedding context. This is the case for regular gratings, but also for homogeneous colour stimuli (56) and redundant, compressible natural scenes (12). Here, natural scenes induced more gamma oscillations than the synthetic stimuli, likely because they contain more compressible features and better match the priors resident in the synaptic weight distribution of cortical networks. However, previous studies on the attentional modulation of gamma oscillations in V1 have reported mixed results (59) and this heterogeneity was reflected also here, across the two monkeys.

One puzzling aspect of our findings is the confinement of the attentional benefits on stimulus encoding to the temporal interval preceding a change in the cued stimulus. Since the task requires only a suppression of reflexive responses to distractor change, no enhancements in encoding for the natural scenes are necessary or even expected. Yet these enhancements in stimulus discriminability occurred close to the anticipated stimulus change and were more pronounced in trials with short reaction times (Figure S1B). It is conceivable that these states are particularly favorable to permit refinement of stimulus representations by structured top-down signals, suggesting that in a different task context they may carry behavioral relevance.

In summary, we showed that the spatial allocation of attention towards a natural stimulus can engage mechanisms that exploit the higher order statistical regularities of natural images, resulting in enhanced decodability of neuronal population responses in area V1. The alignment between the attentional and natural stimulus variance in low-dimensional projections of V1 activity vectors, which was absent for synthetic low-level stimuli, suggests that attention can involve mechanisms optimized for the processing of natural images in order to refine stimulus representation in V1. These results highlight the importance of using natural stimuli when studying sensory processing and provide important insights into how such factors as natural image statistics and the animals’ internal models of the visual world are central to visual processing even at early levels.

## Materials and Methods

### Electrophysiological Recordings

The data was obtained from two adult rhesus macaque monkeys, one male (monkey A) and one female (monkey I), aged 8 and 12 years, respectively, during the time of the study. All experimental procedures were approved by the local authorities (Regierungspräsidium, Hessen, Darmstadt, Germany) and were in accordance with the animal welfare guidelines of the European Union’s Directive 2010/63/EU. Animals were housed in rooms with outdoor access to a play area and had regular veterinary care and balanced nutrition. The recording chamber was implanted under general anesthesia over the primary visual cortex, the exact location was determined based on stereotactic coordinates derived from MRI and CT scans.

Signals were recorded using a chronically implanted microdrive containing 32 independently movable glass-coated tungsten electrodes with impedance between 0.7 and 1.5 MΩ and 1.5 mm inter-electrode distance (SC32; Gray Matter Research (60)), amplified (TDT, PZ2 pre-amplifier) and digitized at a rate of 24.4 kHz. The signals were filtered between 300 and 4,000 Hz and a threshold was set at 4SD above noise level to extract multi-unit activity. LFP signals were obtained by low-pass filtering at 300 Hz and downsampling to 1.5 kHz.

### Behavioral Paradigm

Animals were seated in a custom primate chair at a distance of 64 cm in front of a 477 × 298 mm monitor (Samsung SyncMaster 2233RZ; 120 Hz refresh rate; gamma-corrected). Stimulation protocols were written using MATLAB (MathWorks) and Psychophysics Toolbox. At the start of each recording week, the receptive fields and orientation preferences of the recorded units were mapped with a moving light bar drifting in a randomized sequence in eight different directions.

The two monkeys performed an attention-modulated change detection task. During the task, eye tracking was performed using an infrared-camera eye-control system (ET-49; Thomas Recording). To initiate a trial, the monkey maintained fixation on a white spot (0.1° visual angle) presented in the center of a black screen and pressed a lever. After 500 ms, two identical visual stimuli appeared in an aperture of 2.8–5.1° at a distance of 2.3–3.2° from the fixation point. One of the stimuli covered the receptive fields of the recorded units, which were situated in the right hemifield, the other stimulus was placed at the mirror symmetric site in the left hemifield. After an additional 700 ms, the fixation spot changed color from white to either red or blue, cuing the monkey to covertly direct its attention to one of the two stimuli (red, right hemifield; blue, left hemifield). When the cued image was rotated (20°), the monkey released a lever in a fixed time window (600 ms for monkey A; 900 ms for monkey I) to receive a reward. A break in fixation (fixation window, 1.5° diameter) or an early lever release resulted in the abortion of the trial, which was announced by a tone signal. Data analysis was based on completed correct trials (mean number per session 967 trials for monkey A; 720 trials for monkey I).

### Eye movements

The percentage of saccadic eye movements over the course of the trial is shown in Figure S8A. Breaks in fixation occurred less frequently in the time interval preceding a stimulus change (1700-1900ms, marked in gray) compared to the baseline (−200-0ms, marked in gray). This difference was highly significant (Figure S8B, Wilcoxon signed-rank test, P = 0.00018, n = 18 sessions in 2 monkeys), suggesting that the eyes were very stable in the time interval of interest. Additionally, we found no differences in the frequency of saccadic eye movements between natural and synthetic stimuli (Figure S8C). Finally, we found no differences in the frequency of saccadic eye movements with attention for neither natural scenes (Figure S8D) nor synthetic stimuli (Figure S8E). Taken together, these results suggest that eye movements are unlikely to be strong contributors to the differences observed in our data.

### Visual Stimulus Design

Stimuli were static, black and white images, presented in a square or circular aperture. Within each recording session, all natural scenes and control images had equal luminance and contrast.

Control images were generated in two ways.

1. Filter-scrambling: synthetic stimuli generated from an image model. Filter scrambling was realized by permuting Gabor filter activations elicited by a natural image across the elements of the complete filter bank. The filter bank was composed of a large set of Gabor functions, fitted to the receptive field (RF) characteristics of the recorded neurons. The positions and orientations of the Gabor functions covered the image uniformly, while their size was matched to the RFs of visual cortical neurons recorded at the same eccentricity. An activation variable determined the level of contribution of each particular Gabor function to the image. The synthetic images were generated by linearly combining the activation-scaled Gabor functions. For each synthetic image, the activations of 500–3,000 Gabor functions were sampled from the empirical distribution of Gabor filter responses to a particular natural image. The resulting control images lacked the higher-order structure of natural scenes but matched their low-level statistical properties.
2. Phase-scrambled images. The 2D fast Fourier transform (FFT) of each natural image was computed to obtain a complex magnitude-phase map. The phase values were scrambled by assigning a random value to each element taken from a uniform distribution across the range (− *π, π*). An inverse FFT was then applied to the resulting magnitude-phase maps to produce scrambled versions of the original natural images. These control images lacked the higher-order structure of natural scenes but matched their frequency spectrum. In a second experiment, recorded in one monkey (monkey I), we contrasted synthetic stimuli with and without higher-order structure. These synthetic images were generated similarly to the control images described in (1) and matched the low-level statistical properties of natural scenes. To add higher-order structure, a subset of Gabor functions were arranged in a manner that produced simple contour-like patterns (example of contour synthetic image in Figure 3D).

### Data analysis

Data analysis was performed using custom code in MATLAB (MathWorks) and the Fieldtrip toolbox (61).

We applied non-parametric statistical tests to avoid assumptions about the distributions of the empirical data. Information about sample variables and size is reported in the results section. Critical results and statistics are reported separately for individual animals in the supplementary materials.

### Discriminability index

The *unit d*’, also known as Cohen’s effect size (62), for a pair of stimuli, was calculated as:

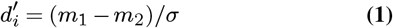

where *m*_1_ and *m*_2_ are the mean spike-counts across trials of unit *i* to the two stimuli and *σ* = (*σ*_1_ +*σ*_2_)*/*2 is the mean standard deviation. This measure was used in Figures 2E, F and S3C and D.

The *population d*’ was calculated similarly, except in this case *m*_1_, *m*_2_ and *σ* are *n*-dimensional vectors of spike-counts, where *n* is the number of simultaneously recorded units in a session. Distances in vector space were calculated using the Euclidean distance. The *population d*’ was used in Figures 1,2, S1 and S2.

### Principal component analysis and stimulus classification

PCA was applied on population spike-count vectors calculated over a 200 ms time window in the trial, 500-700 ms after stimulus onset (pre-cue) and included trials from both attentional conditions. The control stimuli and contour stimuli were analyzed separately, in a similar manner. Thus, the projection space obtained via PCA was different for natural scenes and control stimuli. A comparison of the percentage of variance explained by an increasing number of principal components, for natural scenes and controls recorded in the same sessions, can be found in Figure S6.

To test alignement between the stimulus and the attentional variance (Figure 3), spike-counts over the 1700-1900 ms interval (post-cue) were projected into the PCA space constructed pre-cue. Stimulus classification was performed based on projected data, separately, for trials belonging to the attention in/out conditions. For each attentional condition, Naïve Bayes classifiers were trained to decode the stimulus identity based on data points mapped in the space described by the first *n* principal components, with *n* = 1 to 16. Cross-validation was performed by randomly subsampling the data (*k* 1− data partitions used for training, 1 used for test, *k* repetitions; *k* = 5). This meant that, for each pair of stimuli and each *n*, we ran classifiers *k* = 5 times, on each iteration randomly sampling the population response to the two stimuli. The unseen trials were then used to assess test performance. The performance values reported in Figure 3B are mean validation scores pulled across all stimulus pairs and all recording sessions, with shaded areas representing the standard error of the mean. Chance level was 50 %.

To enquire whether stimulus decoders benefit from knowledge on correlated variability across trials, stimulus classification for variable numbers of PCs was compared for shuffled and unshuffled data in Figure S6. In this case, shuffling was performed across trials, within each stimulus condition (i.e. signal correlations were not affected), after the construction of the PCA projection space, but before the training of the Bayesian classifier. Test data was unshuffled, so that the distribution of original spike-counts was not affected.

Three additional projection spaces were considered for the analysis presented in Figure 4. First, a random orthogonal basis space was generated with the same number of dimensions as the original population space. Second, PCA was computed either on spontaneous activity (spike-count vectors over the -200-0 ms window) or early evoked activity (spike-count vectors over the 100-300 ms window). These projections were performed in individual sessions and averaged over sessions.

### LFP analysis

Power spectra were computed using a frequency-dependent window length (5 cycles per time window). This approach decreases the temporal smoothing at higher frequencies and increases the sensitivity to brief effects. The time-windows were moved in steps of 10 ms and Hann-tapered to avoid spectral leakage.

LFP power differences between trials in the attention-in and attention-out conditions are captured in Figure 1D (18 sessions, precue baseline substracted). A more extensive analysis of attentional differences and stimulus-type differences was performed in individual animals (Figure S4).

Stimulus classification (Naïve Bayes) based on LFP power in various frequency bands, was applied in individual animals, following a similar cross-validation procedure as the one described above for the PCA projections (Figure S7).

## Acknowledgments

We thank Gareth Bland for providing data management support and technical support in the laboratory. Many thanks to Patrick Jendritza for insightful discussions in the early days of the project. This work was supported by the Human Frontier Science Program (RGP0044/2018, AL, GO and WS), the Reinhart Kosselleck grant of the German Research Foundation (WS) and the European Union project RRF-2.3.1-21-2022-00004 within the framework of the Artificial Intelligence National Laboratory (GO).

## Supplementary Materials

**Fig. S1.**
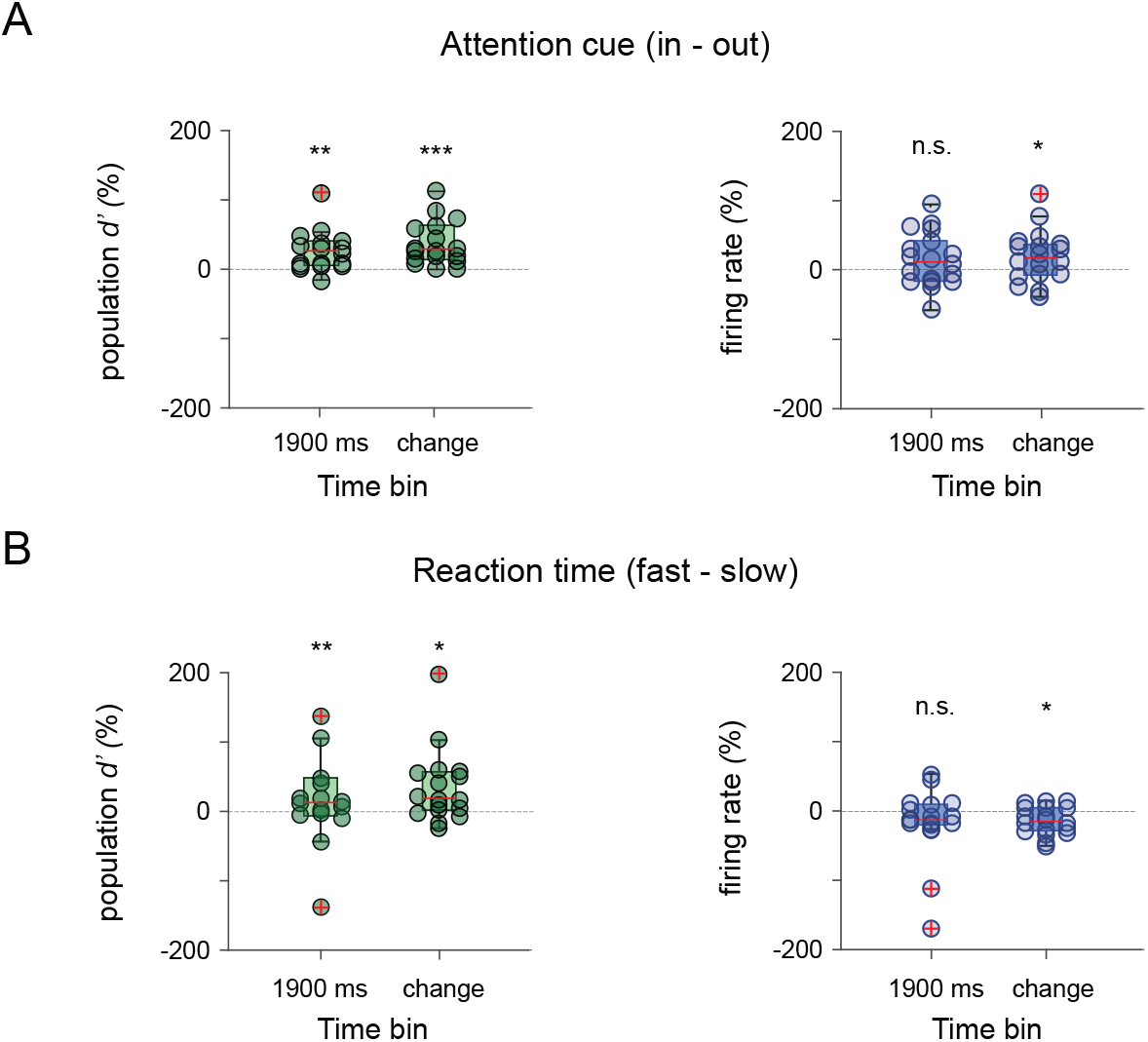
Attentional changes in neuronal population responses. A) Population discriminability index *d*’ (left) and population firing rates (right) shown as percentage increase between the *attention-in* and *attention-out* conditions. Circles represent independent recording sessions. The box-plots correspond to two time windows in the trial (1700-1900 ms and the 200 ms window before the stimulus change). Changes in population *d*’ with attention were strongly significant in both windows (Wilcoxon signed-rank test; 1700-1900 ms: P = 0.0012; change-aligned: P = 0.00023; n = 18 sessions). Firing rates increased significantly for the change-aligned data (Wilcoxon signed-rank test; 1700-1900 ms: P = 0.1330 *n*.*s*.; change-aligned: P = 0.0279; n = 18 sessions). B) Comparison of population *d*’ and population firing rates as a function of reaction time (RT) to the stimulus change. Trials corresponding to the *attention-in* condition, were sorted based on RT, separately for each stimulus condition, and split in two halves referred to as *fast-RT* and *slow-RT*. We found that the population discriminability index was higher for fast RT compared to slow RT (Wilcoxon signed-rank test; 1700-1900 ms: P = 0.0043 ; change-aligned: P = 0.0123; n = 18 sessions). Population firing rates were significantly lower for the change-aligned data (Wilcoxon signed-rank test; 1700-1900 ms: P = 0.6791 *n*.*s*.. ; change-aligned: P = 0.0347; n = 18 sessions). Note that the results from the subplots in B were based on approximately half the number of trials from the subplots in A.

**Fig. S2.**
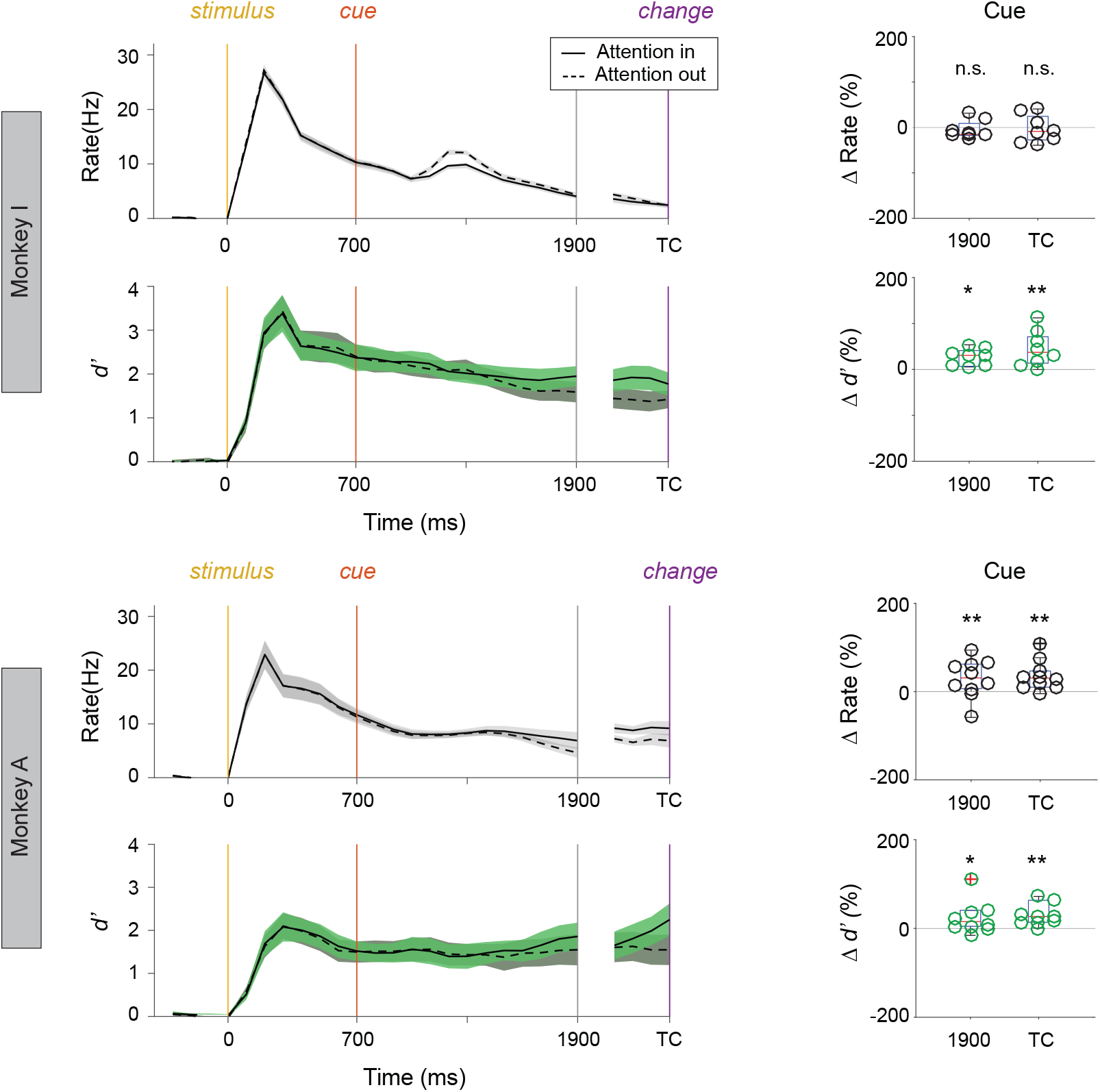
Firing rate responses and discriminability index *d*’ in individual animals. Responses are shown over the course of the trial: spike-counts were calculated over 200 ms windows with 100 ms sliding resolution. Population firing rates (blue) increased significantly with attention, before the stimulus change, in one monkey (right panels, blue; Wilcoxon signed-rank test; 1700-1900 ms: P = 0.46 n.s.; change-aligned: P = 0.64 n.s.; n = 8 sessions Monkey I; 1700-1900 ms: P = 0.009; change-aligned: P = 0.0039; n = 10 sessions Monkey A). Population *d*′ (green) increased significantly with attention, before the stimulus change in both monkeys (right panels, green; Wilcoxon signed-rank test; 1700-1900 ms: P = 0.0078; change-aligned: P = 0.0156; n = 8 sessions Monkey I; 1700-1900 ms: P = 0.04; change-aligned: P = 0.002; n = 10 sessions Monkey A). Shading marks standard error of the mean. Right panels show changes with attention in all individual recording sessions for data aligned on both time of stimulus-onset and stimulus-change.

**Fig. S3.**
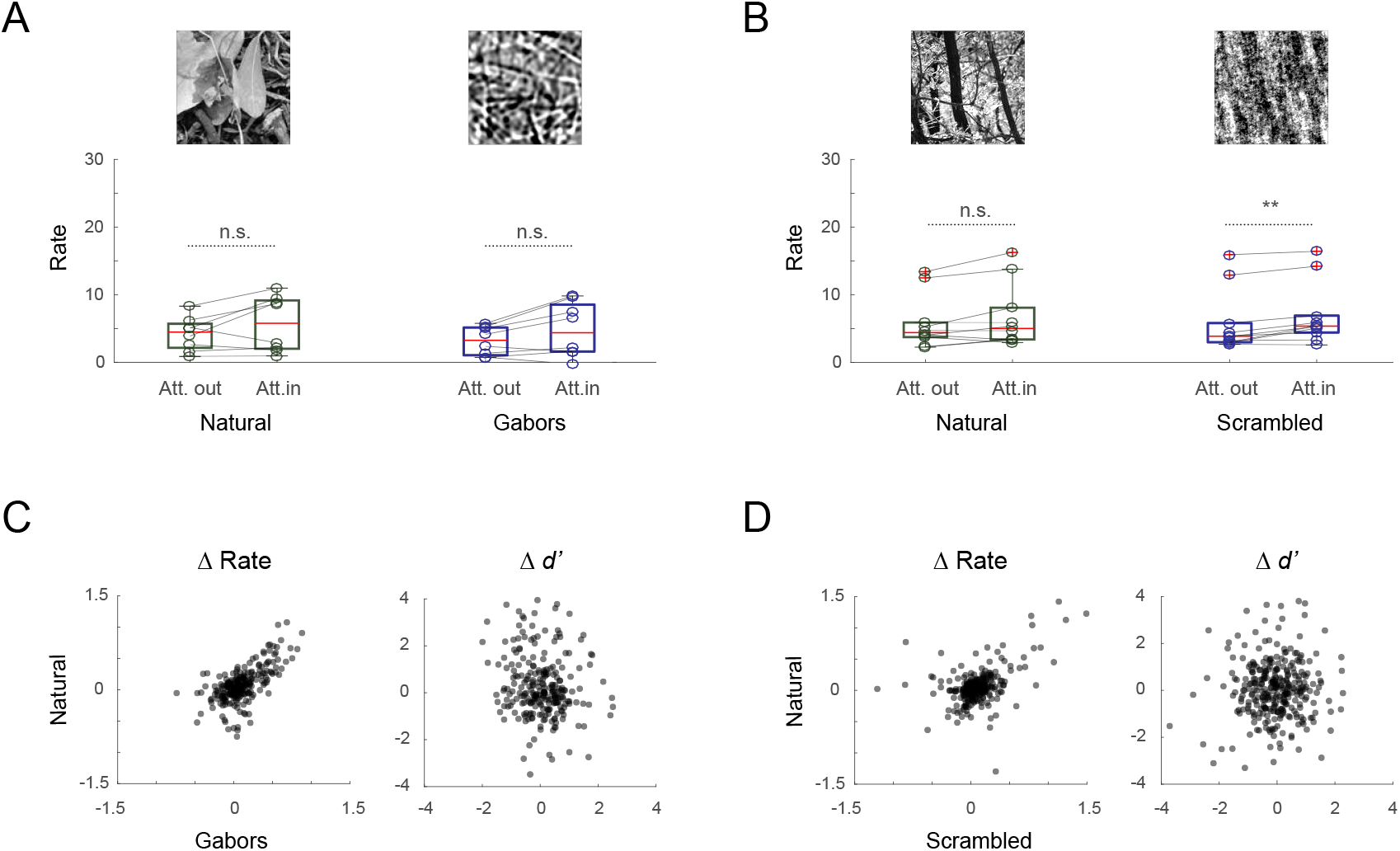
Firing rate responses are modulated by attention similarly for natural scenes and synthetic controls. (A) Firing rate responses for both natural scenes and synthetic gabor-images increase modestly with attention, however the increase is not significant (Wilcoxon signed-rank test; change-aligned; natural-scenes P = 0.1484; gabors P = 0.0781; n = 8 sessions, 2 monkeys). (B) Firing rate responses for natural scenes and scrambled images increase modestly with attention. The increase is only significant for the scrambled images (Wilcoxon signed-rank test; change-aligned; natural-scenes P = 0.084 n.s.; scrambled P = 0.0039; n = 10 sessions, 2 monkeys). C) Firing rate changes with attention for natural scenes and synthetic gabor-images are positively correlated across units (values are z-scored per session); changes in *d*′ are not (Spearman’s rank correlation; firing rate *r*= 0.58, *p* = 2.4e-24; *d*′ *r*= -0.16, *p* = 0.009). D) As in C, firing rate changes with attention for natural scenes and scrambled scenes are positively correlated across units; changes in *d*′ are not (Spearman’s rank correlation; firing rate *r*= 0.35, *p* = 1.1e-10; *d*′ *r*= 0.009, *p* = 0.87).

**Fig. S4.**
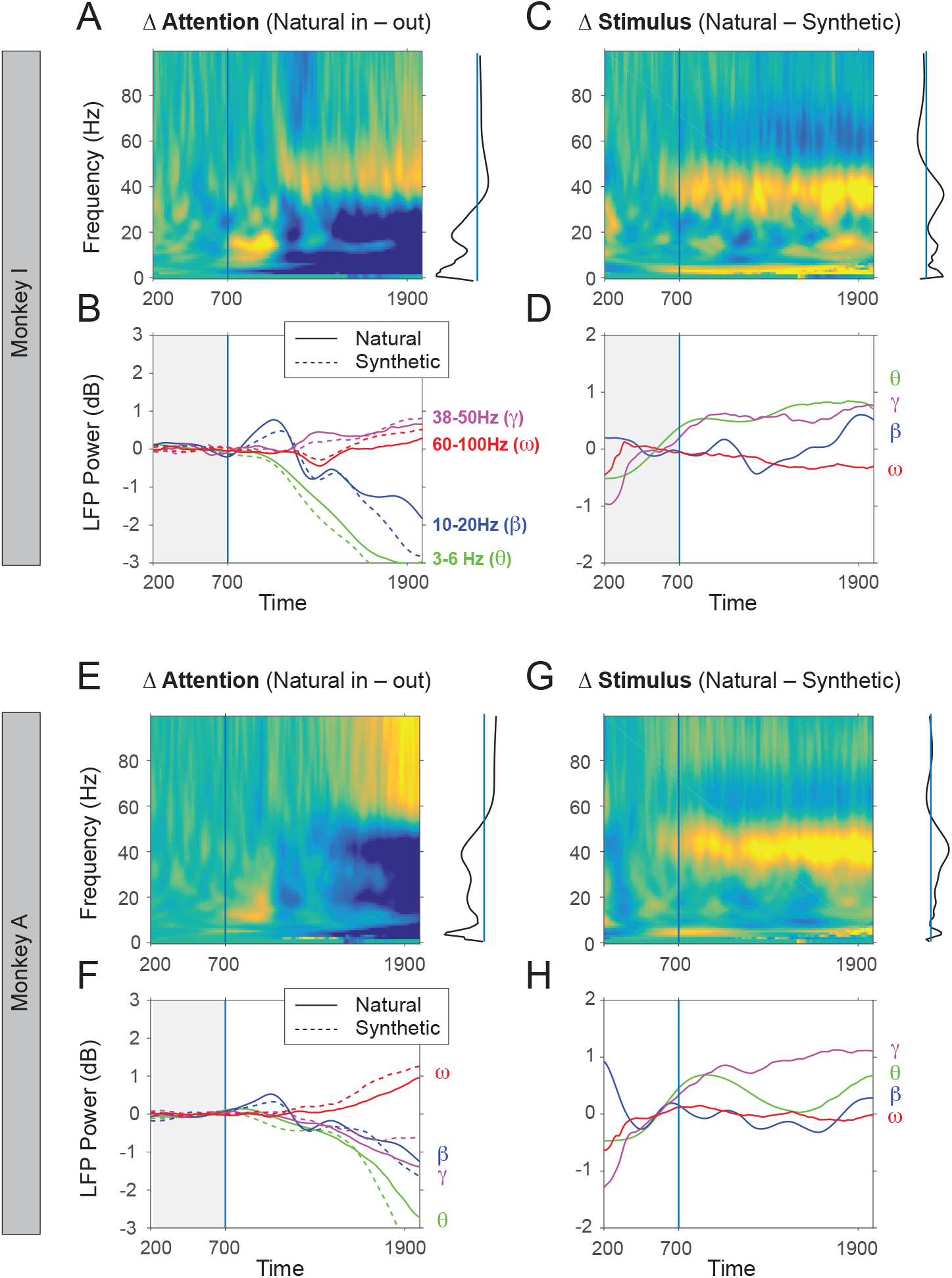
Trial dynamics of LFP power spectra in two monkeys (upper and lower panels) (A) Attentional differences in time-frequency log-transformed LFP power in monkey I (8 recording sessions, 500 ms pre-cue baseline was substracted). The arrival of the attentional cue at 700 ms is marked by vertical line. Right-side projection shows attentional modulation of LFP power for the 1700-1900 ms time window. (B) Attentional differences in LFP power in 4 frequency bands for natural stimuli (continuous lines) and synthetic stimuli (dotted lines). In this monkey, the LFP power at frequencies >38Hz increased with attention, while the LFP power at frequencies <38Hz decreased with attention. (C) Differences in LFP power between natural scenes and synthetic images for the attention-in condition. 500 ms pre-cue baseline was substracted to emphasize post-cue effects. (D) Stimulus differences in LFP power in the gamma (38-50Hz) and theta (3-6Hz) ranges were stronger for natural stimuli compared to their synthetic counterparts. (E) Attentional differences in time-frequency log-transformed LFP power for monkey A (8 recording sessions). In this monkey, the LFP power at frequencies >50Hz increased with attention, while the LFP power at frequencies <50Hz decreased with attention. (F) Attentional differences in LFP power in 4 bands for monkey A. (F) Differences in LFP power between natural scenes and synthetic images for the attention-in condition for monkey A (8 recording sessions). (G) Stimulus differences in 4 bands. Similarly to the plots in (D), natural stimuli produced more gamma and more theta compared to the synthetic controls.

**Fig. S5.**
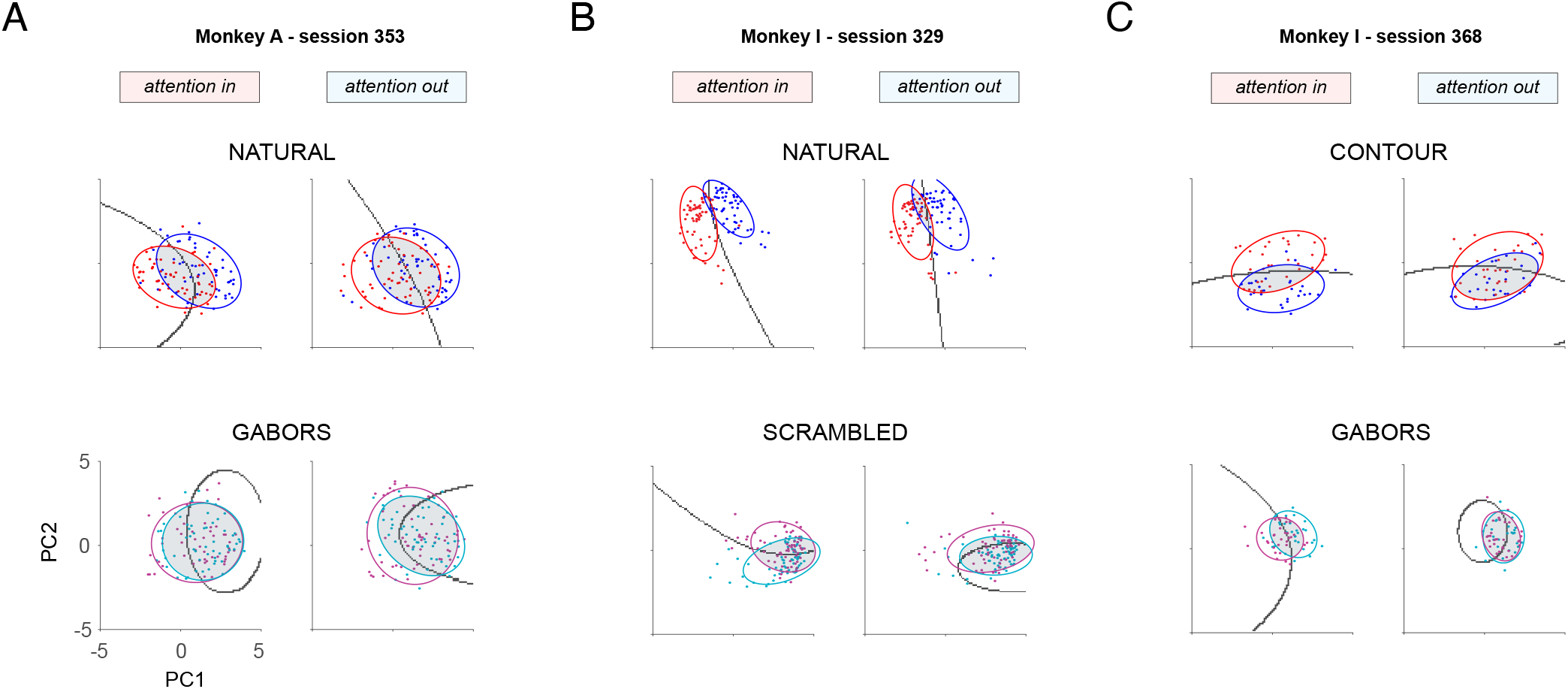
Effects of attention on stimulus encoding in principal component space. Three example sessions (A, B and C), contrasting pairs of natural stimuli and contour stimuli (upper panels), to synthetic controls (bottom panels). Individual points represent trials, each point is a spike-count vector over the 1700-1900 ms interval projected in PCA space. As in Figure 3, the PCA space was constructed based on pre-cue activity (500-700 ms). Ellipses were fit to encompass responses within one standard deviation from the mean. Gray lines show boundaries of Bayesian decoders. For natural scenes and contour stimuli, overlap between the population responses is lower in the *attention-in* condition. Thus the attentional effects can be observed already from a low number of principal components, suggesting an alignment between attentional and stimulus variance. Number PCs

**Fig. S6.**
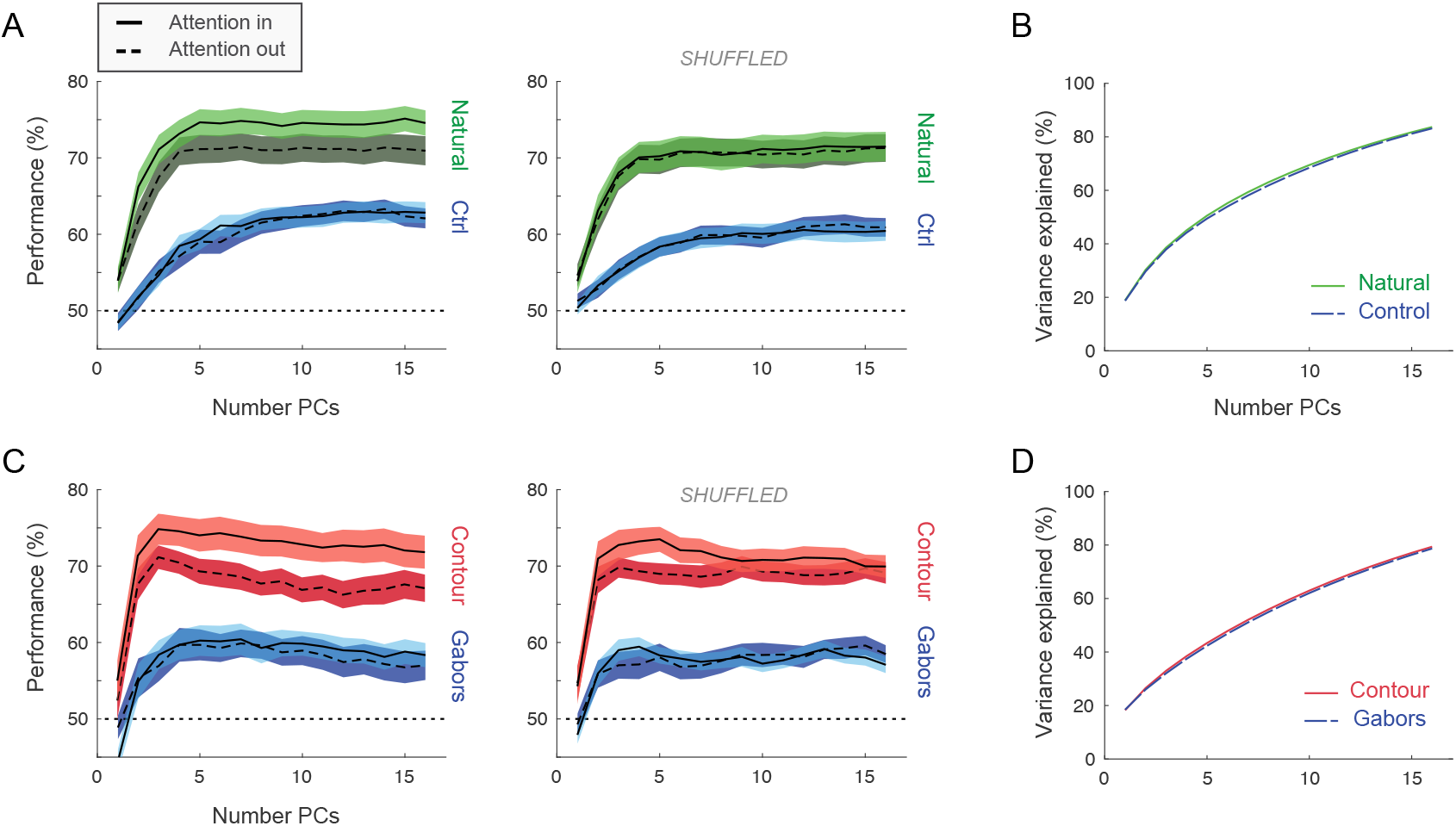
Effect of trial-shuffling on stimulus decoding in PCA space. (A) Decoding performance of natural scenes (green) and control stimuli (blue) based on spike-count vectors over the 1700-1900 ms interval in principal component space, for a variable number of PCs (*x*-axis). As in Figure 3, the PCA space was constructed separately for natural scenes and controls, based on pre-cue activity (500-700 ms, see more details in Methods). Original data (left; n = 71 stimulus pairs; 18 recording sessions) is contrasted to trial-shuffled data (right). Shuffling removes correlations across trials, i.e. the so called noise correlations or spike-count correlations. Shuffling affected the fit of the Bayesian classifier, not the construction of the PCA space, and was performed only on training data, not on test data. Shaded areas indicate standard error of the mean. (B) Variance explained by an increasing number of principal components is similar for PCA spaces constructed separately for natural scenes and controls. These PCA spaces are identical for shuffled data, since shuffling was only applied before training the classifier. (C) Similar to A for contour stimuli and gabor controls (example stimuli in Figure 3) (D) similar to B for contour stimuli and gabor controls (n = 30 stimulus pairs; 5 recording sessions). In A and C, attentional effects can be observed already from a low number of principal components, for natural scenes (green) and contour stimuli (red), but are absent for the synthetic controls (blue) and are reduced by shuffling (right panels). In B and D, variance explained is almost identical for PCA spaces constructed for natural stimuli/contour stimuli and their respective control counterparts Lazar *et al*. | Paying attention to natural scenes in area V1 bioR*χ*iv | 17

**Fig. S7.**
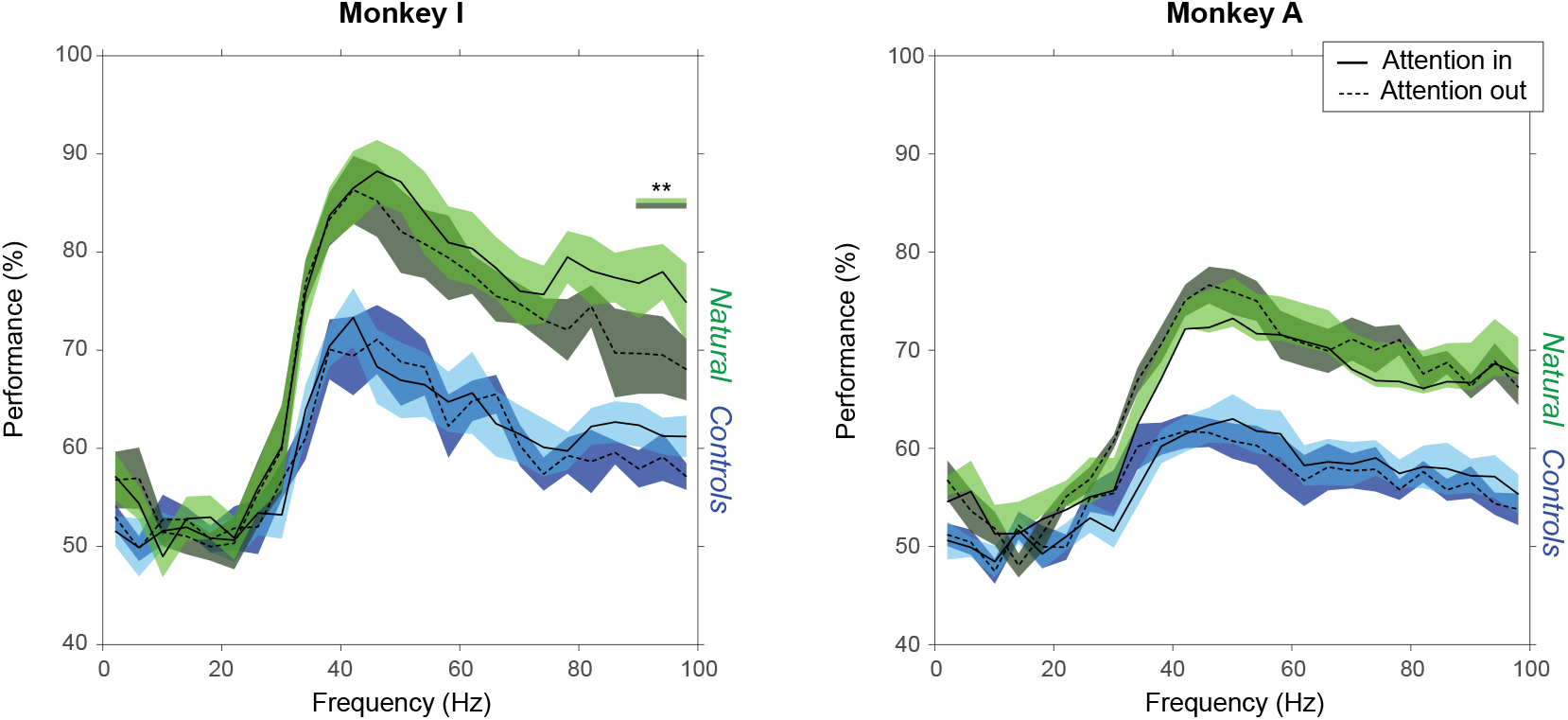
Decoding of stimulus identity based on LFP responses in individual animals. Bayesian classifiers were used to decode the identity of a stimulus based on single trial LFP data (monkey I left, n = 8 sessions, monkey A right, n = 10 sessions; 1700-1900ms time-window). In each recording session the LFP power of all channels (maximum 32) for 4 consecutive frequencies (1-4Hz, 5-8Hz.. 96-100Hz) were concatenated, resulting in data vectors of length < 4×32. Performance is shown separately for natural scenes (green) and control images (blue), for all 25 frequency intervals (x-axis). In both monkeys, classification of natural scenes was significantly above chance level and above the classification performance of control images, for frequencies higher than 40Hz (10-fold validation, chance level 50%). There was no attentional effect on stimulus decoding based on LFP power, except at the highest frequencies in monkey I (88-91Hz, 92-95Hz, 96-100Hz, significant intervals marked by green bar).

**Fig. S8.**
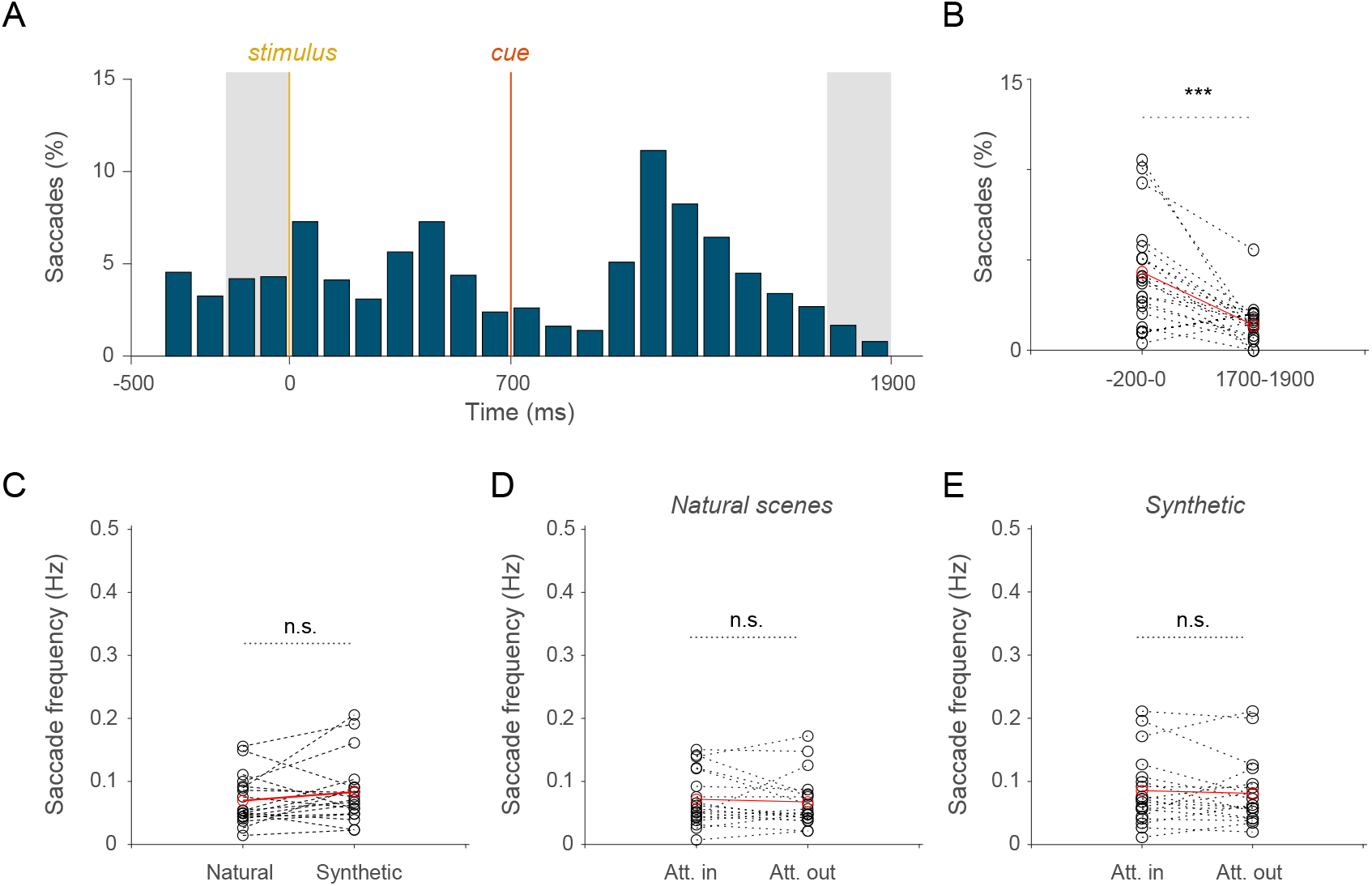
Eye movements during the task. (A) Frequency of saccadic eye movements along the trial. The time window preceding the stimulus change (1700-1900ms, marked in gray), in which the effects of visual attention were strong, corresponds to a period with reduced eye movements compared to the baseline (−200,0ms, marked in gray). (B) Percentage of saccadic eye movements before the stimulus change is shown compared to the baseline across sessions (Wilcoxon signed-rank test,, P = 0.00018, n = 18 sessions in 2 monkeys). (C) Saccades towards natural and synthetic stimuli occurred with similar frequency (left panel; Wilcoxon signed-rank test, post-cue time interval 700-1900ms, P = 0.17 n.s.). (D) Saccades towards attended and unattended natural stimuli occurred with similar frequency (right panel; Wilcoxon signed-rank test; 700-1900ms; P = 0.49 n.s.). (E) Saccades towards attended and unattended synthetic stimuli occurred with similar frequency (right panel; Wilcoxon signed-rank test; 700-1900ms; P =0.50 n.s.).

